# The rostral intralaminar nuclear complex of the thalamus supports striatally-mediated action reinforcement

**DOI:** 10.1101/2022.10.25.513738

**Authors:** Kara K. Cover, Abby G. Lieberman, Morgan M. Heckman, Brian N. Mathur

**Affiliations:** Department of Pharmacology, University of Maryland School of Medicine, Baltimore, MD USA 21201

## Abstract

The dorsal striatum (DS) mediates the selection of actions for reward acquisition necessary for survival. Striatal pathology contributes to several neuropsychiatric conditions, including aberrant selection of actions for specific rewards in addiction. A major source of glutamate driving striatal activity is the rostral intralaminar nuclei of the thalamus (rILN). Yet, the information that is relayed to the striatum to support action selection is unknown. Here we discovered that rILN neurons projecting to the DS integrated information from several subcortical and cortical sources and that these rILN→DS neurons stably signaled at two time points in mice performing an action sequence task reinforced by sucrose reward: action initiation and reward acquisition. *In vivo* activation of this pathway increased the number of successful trials whereas inhibition decreased the number of successful trials. These findings illuminate a role for the rostral intralaminar nuclear complex in reinforcing actions.

## Introduction

Classic models of basal ganglia function regard the dorsal striatum (DS) as an integrator of cortical glutamatergic signaling that is modulated by reward signals arising from midbrain dopaminergic input to support the reinforcement of actions directed toward the acquisition of rewards necessary for survival (Albin et al., 1989; DeLong & Wichmann, 2009; Gerfen & Surmeier, 2011; Kreitzer & Malenka, 2008). While several studies highlight the fact that thalamic nuclei directly project to the striatum (Alloway et al., 2014; J. Ding et al., 2008; Parker et al., 2016; Smith et al., 2014), these nuclei are far less represented in basal ganglia models. This reflects our relatively poor understanding of the functional role of thalamostriatal signaling in action reinforcement.

The intralaminar nuclei are the primary source of thalamic excitatory input to the striatum (Elena Erro et al., 2002; Parent et al., 1983). These nuclei are segregated into rostral and caudal subdivisions. At the caudal end, the medullary lamina splits to create the parafascicular nucleus, and more laterally, the centré median nucleus in monkey and human brains. The boundary distinguishing these two nuclei is undetectable in rodents and other smaller mammals; thus, the posterior intralaminar nuclei are referred to solely as the parafascicular nucleus with the consideration that the lateral component of this nucleus is homologous to the centré median nucleus (Jones, 2007). At the rostral end, the so-called rostral intralaminar nuclei (rILN) include the poorly described and weakly demarcated central lateral (CL), paracentral (PC), and central medial (CM) regions. Both the caudal intralaminar grouping and the rILN receive input from a wide array of cortical and subcortical centers and output to both the striatum and the cortex. As such, it is unclear which of the inputs to the intralaminar nuclei are routed to the striatum to potentially influence action reinforcement (Groenewegen & Berendse, 1994).

Functionally, the caudal intralaminar nuclear group responds to salient sensory cues in monkey (Minamimoto & Kimura, 2002) and, in rodents, detects changes in action-outcome contingencies to facilitate behavioral flexibility (Bradfield et al., 2013; Bradfield & Balleine, 2017; Yamanaka et al., 2018). Recent work implicates the rILN in facilitating the execution and switching of learned actions (Kato et al., 2018) and driving behavioral reinforcement (Cover et al., 2019; Johnson et al., 2020). Together, these data suggest that the rILN contribute to action reinforcement, but the information that the rILN integrates and relays to the striatum to support this function is unknown. Examining this in mice using in vivo neural circuit monitoring and manipulations methods, we discovered that the rILN dynamically signal at both the initiation of an action and during reward acquisition to optimize action performance. We also determined that striatal-projecting rILN neurons demonstrate homogenous physiological properties across the three regions but differently integrate cortical, midbrain, and hindbrain information that is then passed on to the striatum. These data support the notion that the rILN are a central integrator contributing to basal ganglia-mediated action reinforcement.

## Results

### Striatal-projecting rILN neurons exhibit uniform properties

The rILN consist of a contiguous band of cells within the internal medullary lamina that is parcellated into CL, PC, and CM nuclei (Berman & Jones, 1982). Although anatomical tracing suggests subtle differences in afferent and efferent connectivity among these three subregions (Van der Werf et al., 2002), it is unclear whether these nuclei represent functionally distinct cellular groups. To investigate whether rILN→DS neurons exhibited physiological differences across the three nuclear divisions, we injected a retrograde traveling tdTomato-expressing virus in the central DS of mice and assessed neurophysiological properties of rILN→DS neurons using whole-cell patch clamp electrophysiology in acute slices (Fig. 1A, B). We found that neurons across the three nuclei did not significantly differ in membrane capacitance (Fig. 1C; F(2,40)=1.242, P=0.30), membrane resistance (Fig. 1D; F(2,40)=1.212, P=0.31), input resistance (Fig. 1E; F(2,40)=0.520, P=0.60), or resting membrane potential (Fig. 1F; F(2,40)=0.124, P=0.88).

**Figure 1.**
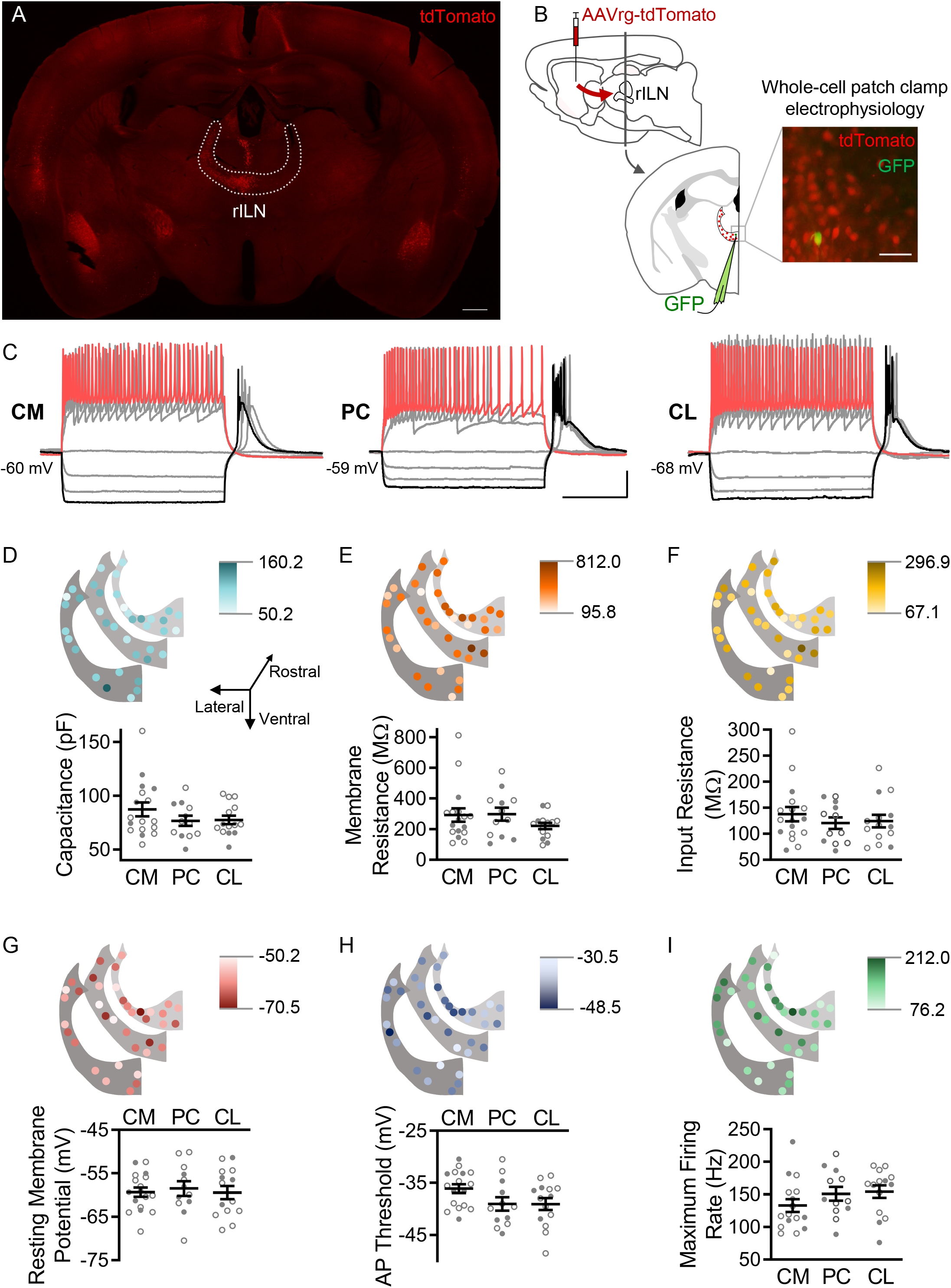
Dorsal striatum-projecting neurons of the rostral intralaminar thalamus (rILN→DS) exhibit uniform physiological properties. A. rILN→DS neurons project ipsilaterally. Coronal mouse brain section through the rILN. Following AAVrg-tdTomato injection in the left dorsal striatum (DS) left rILN→DS neurons labeled with tdTomato. B. Left: Schematic of experimental strategy to label rILN→DS neurons. Right: a patched neuron (GFP) among tdTomato-labeled rILN neurons. C. Representative traces showing central medial (CM), paracentral (PC), and central lateral (CL) neuronal responses to 500 ms current injection steps. D. Top: Neuronal membrane capacitance of rILN neurons mapped across three rostral-caudal coronal planes in light to dark color gradient spanning minimum to maximum capacitance values. Bottom: Membrane capacitance did not differ between CM (n=17), PC (n=12), and CL (n=13) nuclei. E-I. Analysis of membrane resistance (E), input resistance (F), resting membrane potential (G), action potential (AP) threshold (H), and maximum firing rate elicited by current steps (I; CM n=16), for which no significant differences existed between the three rILN nuclei. Scale bars: 500 um (A), 100 μm (B), 20 mV and 200 ms (C). One-way ANOVA (D-I). n = number of cells. Filled and open data points represent cells from male and female mice, respectively.

To assess differences in intrinsic firing properties, we injected a current ramp to induce action potential (AP) firing. We did not observe a significant difference in the AP threshold between the three nuclei (Fig. 1G; F(2,40)=2.684, P=0.08). Following a series of 0.5 s current steps, the maximum firing rate, as calculated from the first six APs, did not differ between neurons from the three rILN nuclei (Fig. 1H; F(2,39)=1.411, P=0.26). AP accommodation was also not different between the nuclei (Fig. S1A; F(2,35)=1.671, P=0.20). We observed that hyperpolarizing current steps induced post-hyperpolarization burst firing in all three nuclei (as shown in Fig. 1B). We did not find differences in either the post-hyperpolarization burst firing inter-spike interval (Fig. S1B; F(2,35)=1.003, P=0.38) or AP number (Fig. S1C; F(2,40)=2.837, P=0.07).

### rILN→DS neurons signal at action initiation and reward acquisition

Establishing that rILN→DS neurons display homogenous intrinsic electrophysiological properties, we next endeavored to elucidate when this pathway is active during action learning and performance. We fiber photometrically recorded activity-dependent calcium signaling selectively from rILN→DS neurons in mice learning a lever pressing operant task (Fig. 2A). To assess whether rILN→DS neurons respond to sensory stimuli eliciting an action, action initiation, termination, kinematic properties, or reward, we trained mice to respond to the lever extension by pressing five times in a defined period of time for a sucrose pellet reward (Fig. 2B). In early training stages mice must complete one, three, or five fixed-ratio presses (FR1, FR3, FR5, respectively) with no time limit (NTL) for completion. Intermediate schedules consist of progressively shortened response periods to complete the FR5, terminating at a 5 s time limit. We found transient increases in rILN→DS activity that accompanied the first press in the action sequence across all stages of training (Fig. 2C). Comparing the average rILN→DS activity z-score at 1.5 s before and at the time of the first press, we observed significant differences from the first training protocol (fixed ratio 1; FR1), an intermediate schedule (FR5-NTL), and the final training schedule (FR5-5s time limit; time relative to press: F(1,12)=26.73, P=0.0002; schedule: F(2,24)=9.587, P=0.0009). We observed similar positive fluctuations in rILN→DS activity aligned to subsequent presses in early training protocols in which mice complete the FR lever-pressing task over an extended period (Fig. 2D-F). This activity was not observed for subsequent presses in mice on the terminal training protocol, however (Fig. 2D-F; schedule: F(1,12)=34.79, P<0.0001; press: F(3,36)=2.146, P=0.11).

**Figure 2.**
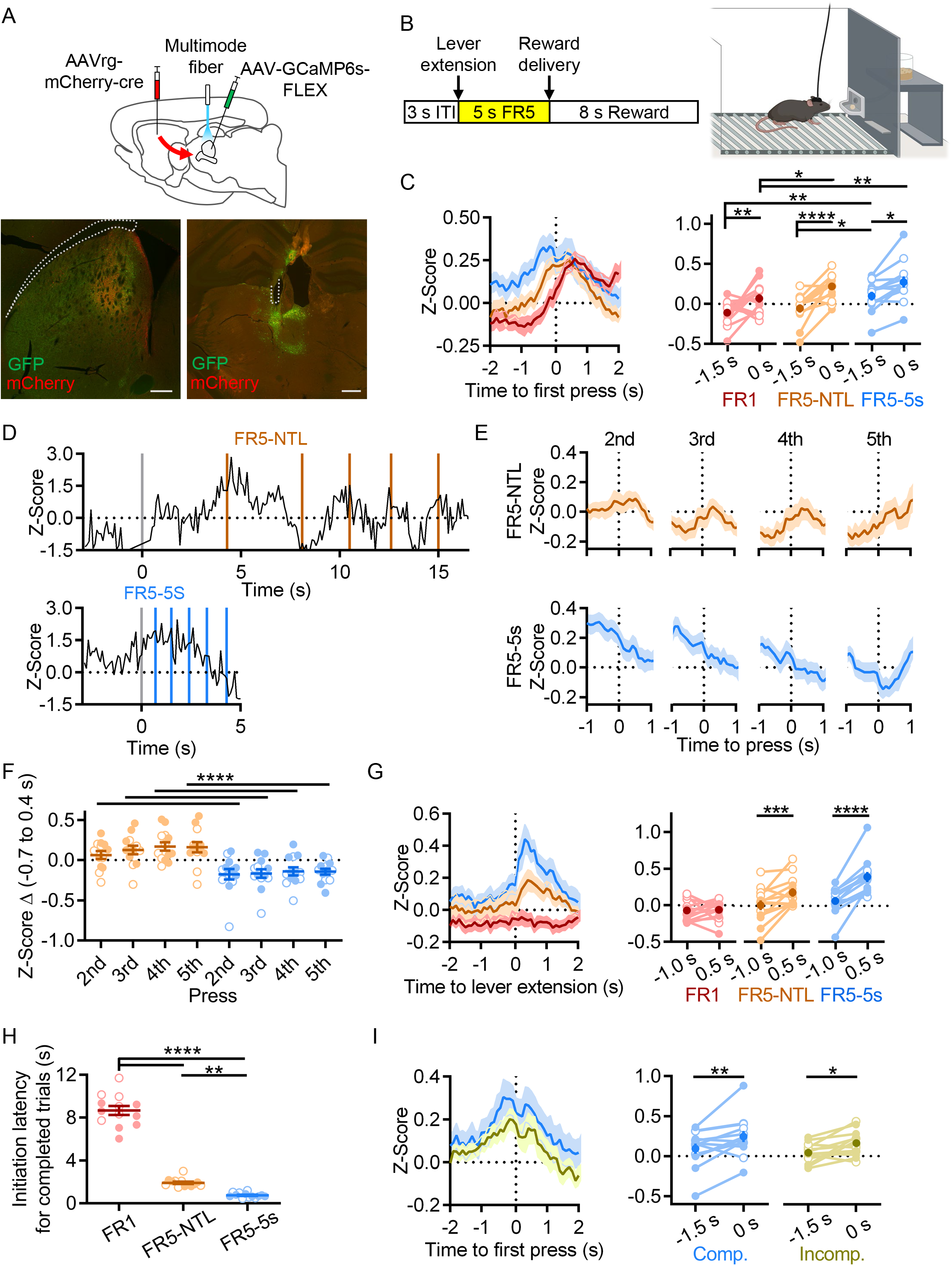
rILN→DS activity aligns to action initiation. A. Top: Experimental strategy for fiber photometric recording of rILN→DS neurons. Bottom: Representative expression of cre recombinase (mCherry) and cre-dependent GCaMP6s (GFP) in the DS (left) and rILN (right) with fiber placement in the rILN (right). B. Left: Schematic of lever press trial design. Each trial starts with a 3 s inter-trial interval (ITI) followed by lever extension. A sucrose pellet is delivered following the fifth press, with an 8 s consumption period before the next trial. Right: Cartoon of operant chamber with lever and sucrose pellet dispenser. C. Left: Average calcium-dependent activity of rILN→DS neurons aligned to the first press for the first training schedule (red; Fixed-rate 1 - no time limit; FR1-NTL), an intermediate protocol (orange; FR5-NTL), and the terminal training schedule (blue; FR5-5s time limit). Right: rILN→DS activity was significantly greater at the time of the first press compared to 1.5 s prior for all three operant schedules (N=13). D. Representative rILN→DS activity from a mouse completing the FR5 sequence on FR5-NTL (top) and FR5-5s (bottom left) schedules. Gray lines indicate time of lever extension and colored lines show individual lever presses. E. Average rILN→DS activity for presses 2-5 from FR5-NTL (top) and FR5-5s (bottom) schedules. F. The change in z-score from −0.7 s to 0.4 s (relative to press) was positive for all presses on the FR5-NTL schedule (orange) and negative for all presses on the FR5-5s schedule. G. Left: Photometrically recorded rILN→DS activity aligned to the extension of the lever on completed FR5 trials from FR1 (red), FR5-NTL (orange), and FR5-5s (blue) training schedules. Right: rILN→DS signaling increased following lever extension on all schedules except for FR1. H. The average latency to initiate pressing progressively decreased with training. I. Left: first press-aligned rILN→DS activity from completed (blue) and incomplete (yellow) FR5 trials in trained mice. Right: The average rILN→DS activity change from −1.5 to 0 s relative to first press differed by trial performance. Scale bars: 500 μm. Two-way repeated measures ANOVA (C, F, G, I); one-way repeated measures ANOVA (H). N = number of mice. Filled and open data points represent male and female mice, respectively.

We next examined how rILN→DS neurons activate relative to specific elements of the FR5 task. We found that rILN→DS activity peaked following the extension of the lever into the operant chamber on completed trials in intermediate and terminal stages of training and that the magnitude of this signal change increased with training (Fig. 2G; time relative to lever: F(1,12)=17.27, P=0.0013; schedule: F(3,36)=14.49, P<0.0001; interaction: F(3,36)=6.776, P=0.001). The average latency to initiate the lever press for completed trials was 8.7 s for FR1 and decreased to 1.9 seconds (FR5-NTL), and 0.76 s (FR5-5s) (Fig. 2H; F(1.218,14.61)=303.0, P<0.0001). Gross rILN→DS activity at this time, as measured by area under the photometric signal, was negatively correlated with initiation latency on completed trials across all FR5 schedules (r=-0.161; −0.185 and −0.137 95% CI, P<0.0001, n=6338 trials). Thus, we examined whether this lever press-associated rILN→DS signal varied by task performance. We found that a larger signal accompanied completed trials as compared to incomplete trials (in which mice pressed 1-4 times) (Fig. 2I; time relative to first press: F(1,12)=14.73, P=0.0024; trial type: F(1,12)=2.374, P=0.15).

To further explore this rILN→DS signal, we investigated whether rILN→DS activity correlates with general movement by recording mice freely moving in an open arena (Fig. S2A). We observed increases in rILN→DS signaling aligned to movement onset (Fig. S2B; t=2.817, P=0.037) and preceding the maximum velocity achieved during movement epochs (Fig. S2C; t=4.116, P=0.009). Significant changes in rILN→DS activity did not accompany movement cessation (Fig. S2D; t=0.341, P=0.75).

To discriminate between action initiation and the sensory cues (e.g., lever extension) that may signal action initiation, we used a Pavlovian appetitive conditioning paradigm in which sucrose pellets were administered at the termination of an auditory tone (Fig. S3A-B). We did not find tone-related changes in rILN→DS signaling nor did we observe learning-dependent changes in either the tone-paired group (Fig. S3C; time relative to tone: F(1,5)=2.046, P=0.21; training stage: F(1,5)=2.466, P=0.18) or the tone non-paired control cohort (Fig. S3D; time relative to tone: F(1,5)=2.312, P=0.19; training stage: F(1,5)=0.204, P=0.67).

Alternatively, our observation of training-dependent increases in action initiation-associated rILN activity may reflect the learned expectation of a rewarded action. To test this, we recorded rILN→DS activity after switching FR5 reinforcement to 0%. Mice extinguished FR5 lever pressing within several sessions (Fig. S3F, t=11.69, P<0.0001), but the first press-aligned signal on completed (but unreinforced) FR5 trials did not differ significantly from reinforced sessions (Fig. S3G; time relative to lever: F(1,11)=13.36, P=0.0069; reinforcement: F(1,11)=0.1628, P=0.73). In an alternative paradigm, FR5-trained mice alternated between sessions in which completed trials were reinforced at 100% or 50% probability. Lowering the probability of reinforcement did not alter performance (Fig. S3H; t=0.788, P=0.45) nor did it produce significant differences in first press-aligned rILN activity (Fig. S3I; time relative to press: F(1,10)=5.571, P=0.04; reinforcement rate: F(1,10)=0.304, P=0.59).

Lastly, we observed that rILN→DS activity increases at reward acquisition. Sucrose pellet retrieval was accompanied by increased rILN activity on all FR training protocols (Fig. 3A-B; time relative to retrieval: F(1,12)=57.24, P<0.0001; schedule: F(1,12)=10.42, P=0.007; interaction: F(1,12)=5.311, P=0.040). Additionally, this signal negatively correlated with reward retrieval latency across all FR5 schedules (Fig. 3C; r=-0.123, −0.147 and −0.097 95% CI, P<0.0001). Reward delivery was necessary for this activity, as unreinforced FR5 trials did not elicit changes in rILN→DS activity (Fig. 3D; time relative to head entry: F(1,12)=21.01, P=0.006; reinforcement: F(1,12)=5.610, P=0.035; interaction: F(1,12)=89.14, P<0.0001). On intermediate training schedules, mice frequently check the pellet receptacle in between individual presses. We examined rILN→DS activity during these “premature” receptacle head entries on trials that were ultimately completed and observed that these events were accompanied by possible negative fluctuations in rILN activity (Fig. 3E; time relative to head entry: F(1,12)=3.536, P=0.085; schedule: F(2,24)=0.502, P=0.61; interaction: F(2,24)=0.019).

**Figure 3.**
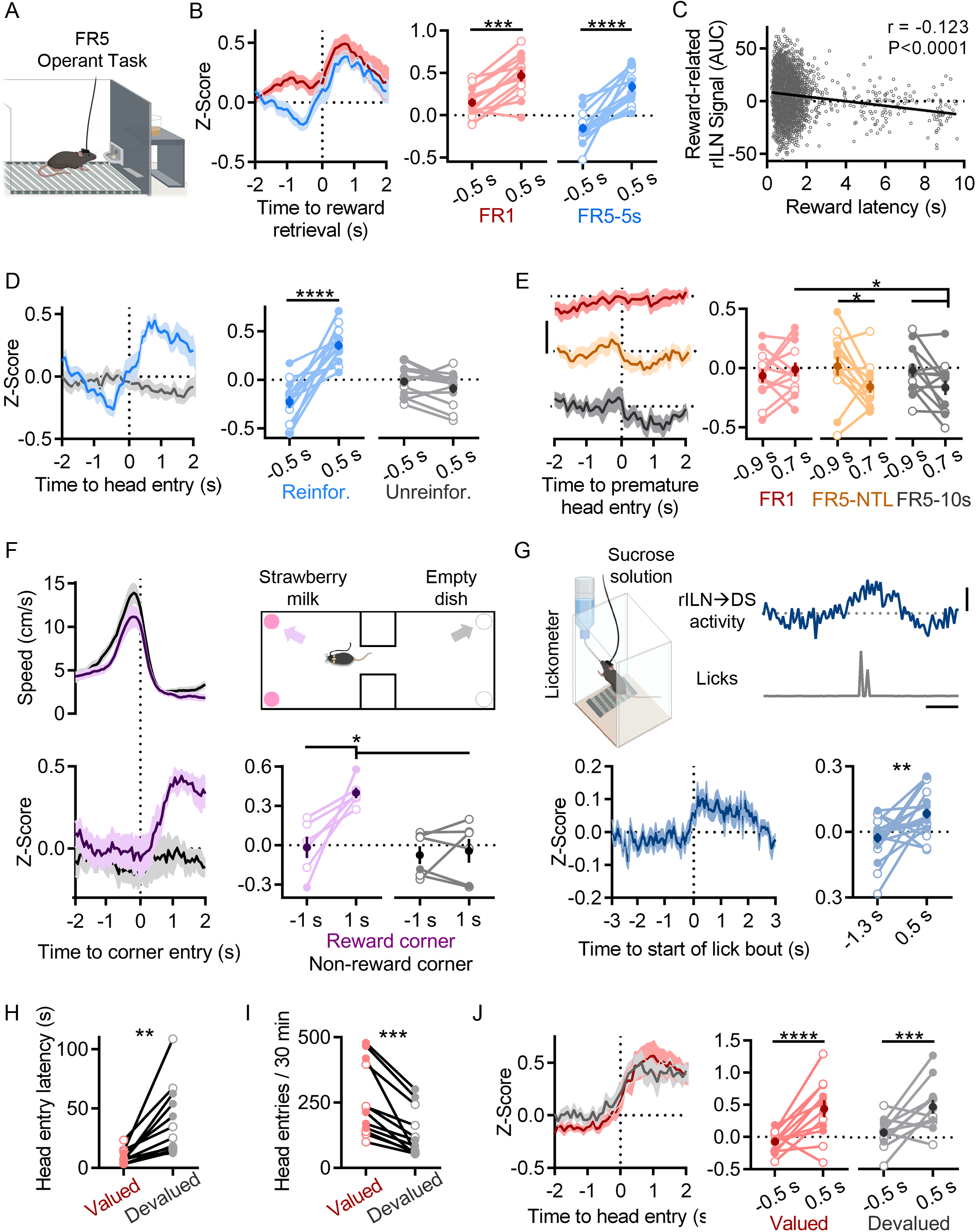
rILN→DS neurons activate with reward acquisition. A. Cartoon of operant chamber. B. Left: rILN→DS activity relative to sucrose pellet retrieval on FR1 (red) and FR5-5s (blue) sessions. Right: rILN→DS activity significantly increased following reward acquisition (N=13). C. Across all FR5 training schedules, reward retrieval-associated rILN→DS activity negatively correlated with retrieval latency (n=6338). D. Left: rILN→DS activity aligned to reward receptacle head entry following completed FR5 trials on reinforced (blue) and unreinforced (gray) FR5-5s sessions. Right: rILN→DS activity increased on reinforced sessions only. E. Left: rILN→DS activity aligned to premature reward receptacle head entries on completed FR1 trials (red), FR5-NTL (orange), and FR5-10s (gray) schedules. Right: rILN→DS activity significantly decreased on FR5-NTL and −10s protocols. F. Movement speed (top left) and corresponding rILN→DS activity (bottom) from mice approaching strawberry milk – containing (purple) and empty (gray) corners of a two-chambered arena (top right) (N=6). rILN→DS activity increased following entry to strawberry milk-baited corners (top right). G. rILN→DS activity relative to sucrose water consumption. Top left: sample trace of rILN→DS activity (blue) and corresponding licks (gray). Top right: Cartoon of drinking chamber. Bottom: average rILN→DS activity aligned to onset of lick bouts (left) which significantly increased following bout onset (right; N=19). H-J. rILN→DS activity was recorded from food-restricted mice over multiple sessions in which sucrose pellets were pseudo-randomly dispensed (valued condition; red). Mice were then recorded for multiple sessions that were preceded by 30-minute free feeding of sucrose pellets in their home cages (devalued condition; gray) (N=12). H. Mice retrieved dispensed sucrose pellets at a slower latency in the devalued state. I. Mice entered the food receptacle fewer times during devalued sessions. J. rILN→DS activity aligned to sucrose pellet retrieval was not significantly different between valued and devalued states. Scale bars: 0.3 z-score (E); 1 s and 0.2 z-score (G). Two-way repeated measures ANOVA (B, D-F, J); linear correlation (C); paired t-test (G, H, I). N = number of mice; n = number of trials. Filled and open data points represent male and female mice, respectively.

We next assessed the generalizability of this reward-related signal in non-operant paradigms. We recorded rILN activity from mice freely moving in an arena with strawberry milk located in two corners. rILN→DS activity increased when mice approached the strawberry milk-baited corners but not the opposing non-baited arena corners (Fig. 3F; time relative to approach: F(1,5)=12.13, P=0.018; corner: F(1,5)=12.54, P=0.017; interaction: F(1,5)=7.965, P=0.037). To directly examine rILN→DS activity relative to reward consumption, we recorded from mice drinking sucrose water from bottles connected to a lickometer. rILN→DS activity significantly increased relative to lick bout onset (Fig. 3G; t=3.210; P=0.005).

We next tested whether reward devaluation may influence rILN→DS activity. Mice were given free access to sucrose pellets prior to 30-minute test sessions in which pellets were pseudo-randomly delivered. Devaluation reduced both latency (Fig. 3H; t=3.953, P=0.002) and frequency of pellet receptable head entries (Fig. 3I; t=5.068, P=0.0004), but did not significantly alter rILN→DS activity aligned to reward retrieval (Fig. 3J; time relative to head entry: F(1,11)=12.23, P=0.003); reward value: F(1,11)=0.642, P=0.44).

### rILN→DS neuronal activity is necessary and sufficient for optimal action performance

To causally test the role of rILN→DS activity, we optogenetically inhibited dorsal striatal projecting rILN neurons in well-trained mice performing the FR5 task (Fig. 4A). Delivering blue light during an epoch encompassing the action initiation and reward retrieval events (White et al., 2018, 2020) pseudo-randomly on 33% of the trials (Fig. 4B) to halorhodopsin-eYFP (NpHR-eYFP) or control eYFP -expressing mice, we found that rILN→DS neuronal inhibition resulted in fewer completed FR5 trials (Fig. 4C; light delivery: F(1,26)=21.90, P<0.0001; virus: F(1,26)=0.390, P=0.54) and more omissions (Fig. 4D; light: F(1,26)=13.99, P=0.0009; virus: F(1,26)=0.127, P=0.72; interaction: F(1,26)=13.10, P=0.0013) (Table S1). We also observed a small but significant effect of light delivery on the percentage of incomplete trials (Fig. 4E; light: F(1,26)=4.834, P=0.037; virus: F(1,26)=2.306, P=0.141).

**Figure 4.**
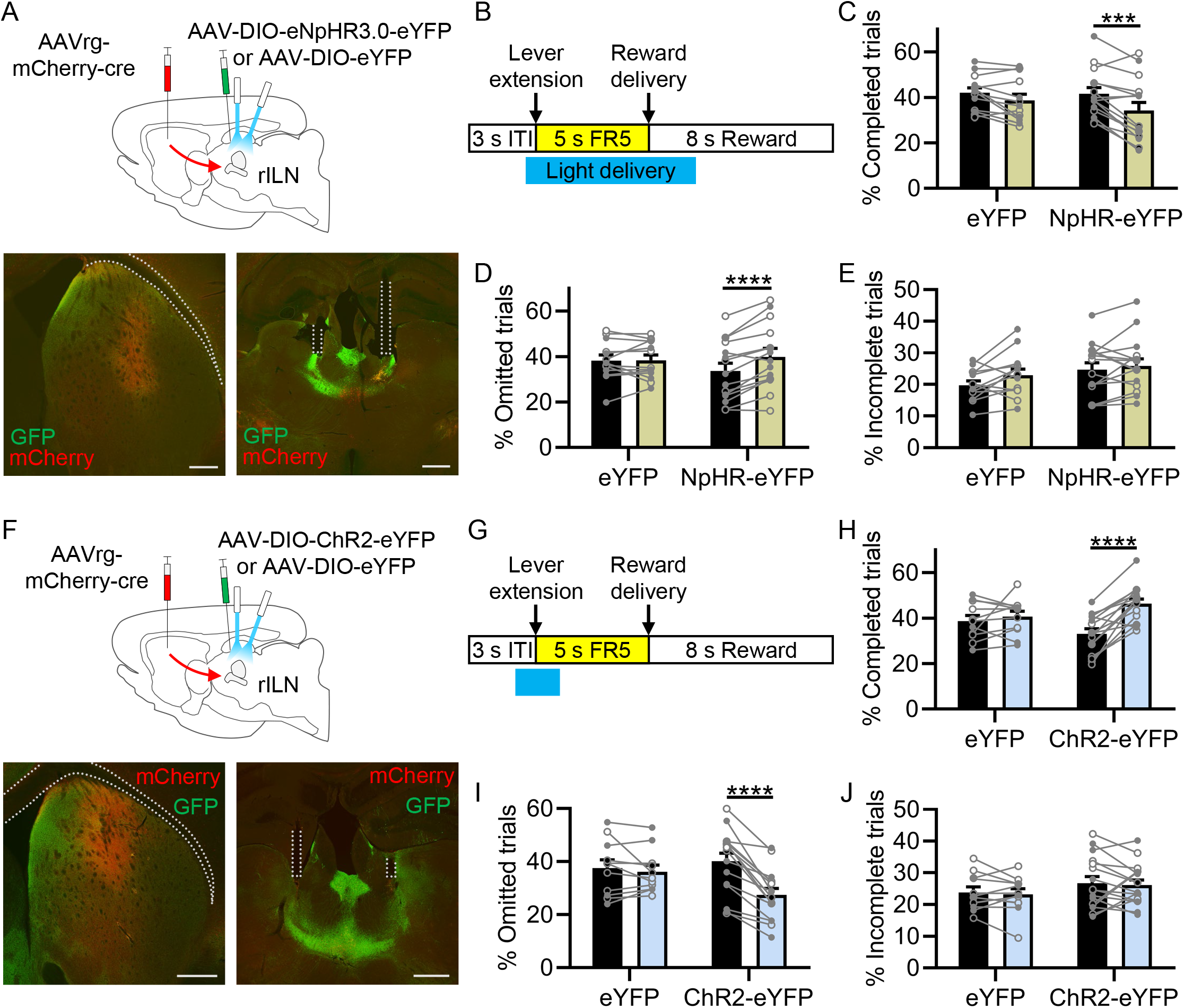
Modulating rILN→DS neuronal activity bidirectionally alters FR5 performance. A. Top: Strategy for optical inhibition of rILN→DS neurons in FR5-trained mice. Bottom: Representative cre recombinase (mCherry) and halorhodopsin (eNpHR-eYFP; GFP) expression in DS (left) and rILN (right) with optical fibers implanted in the rILN. B. Schematic of 470 nm light delivery during the FR5 trial starting 0.5 s prior to lever extension and terminating either 2 s following FR5 completion or at time of lever retraction (for incomplete and omitted FR5 trials). Light was delivered pseudo-randomly on 33% of the trials. C. NpHR-eYFP -expressing (N=15) but not eYFP control (N=13) mice completed fewer FR5s on light-delivered trials (yellow) than non-light-delivered trials (black). D. NpHR-eYFP but not eYFP -expressing mice had a greater omission rate on light-delivered trials. E. Light delivery did not alter the proportion of incomplete FR5 trials for either group. F. Top: strategy for optogenetic activation of rILN→DS neurons. Bottom: representative mCherry-cre (mCherry) and channelrhodopsin (ChR2-eYFP; GFP) expression in DS (left) and rILN (right) with optic fibers implanted in the rILN. G. 2 s 470 nm blue light was delivered 1 s prior to lever extension on 33% of trials. H. ChR2-eYFP mice (N=16) completed more FR5s on light-delivered trials (blue) compared to non-light-delivered trials (black) and eYFP control mice (N=11). I. ChR2-eYFP mice had fewer omissions on light-delivered trials than non-light-delivered trials and eYFP controls. J. There were no significant differences in the rate of incomplete FR5 trials. Scale bars: 500 μm. Two-way repeated measures ANOVA (C-E, H-J). N = number of mice. Filled and open data points represent male and female mice, respectively. See also Table S1.

We previously demonstrated that optogenetic activation of rILN→DS terminals reinforces actions in an intracranial self-stimulation paradigm (Cover et al., 2019). However, it is unknown whether activation of this pathway at action initiation also supports the execution of rewarded action sequences. To test this, we virally expressed channelrhodopsin (ChR2-eYFP) or a control fluorophore (eYFP) in rILN→DS neurons (Fig. 4F). Blue light was delivered on 33% of trials for 2 s starting 1 s prior to lever extension (to promote the lever press action) in trained mice (Fig. 4G). ChR2-eYFP mice completed more FR5s on light-paired trials (Fig. 4H; light: F(1,25)=24.39, P<0.0001; virus: F(1,25)=0.002, P=0.97; interaction: F(1,25)=13.21, P=0.0013) and correspondingly, had fewer omitted trials (Fig. 4I; light: F(1,25)=29.46, P<0.0001; virus: F(1,25)=0.656, P=0.43; interaction: F(1,25)=18.85, P=0.0002), compared to both non-light delivered trials and eYFP controls (Table S1). We did not find significant differences in the number of incomplete FR5 trials (Fig. 4J; light: F(1,25)=0.358, P=0.55; virus: F(1,25)=1.441, P=0.24).

### rILN→DS neurons integrate subcortical and cortical circuits

Although an extensive number of inputs to the rILN are identified (Van der Werf et al., 2002), it is unknown whether these afferents differently synapse on rILN neurons that project to the cortex or striatum. Thus, we sought to qualitatively identify afferents impinging specifically onto rILN→DS neurons. To do this, we applied a viral strategy involving injection of nuclei that are upstream of rILN with an anterograde transsynaptic AAV1 viral construct (AAV1-hSyn-Cre-WPRE-hGH) (Zingg et al., 2017) that delivers cre recombinase to rILN neurons. In addition, we injected a retrogradely traveling cre-dependent tdTomato -expressing virus (AAVrg-CAG-FLEX-tdTomato-WPRE) in the DS. This viral approach produces tdTomato labeling in all rILN→DS neurons postsynaptic to the neurons at the site of AAV1-cre injection (Fig. S4). We used this method to interrogate known excitatory inputs to the rILN. We found that both the anterior cingulate (Fig. 5A) and orbitofrontal (Fig. 5B) cortices synapse on ipsilateral rILN→DS neurons and lesser so on contralateral rILN→DS neurons. We also found that projections arising from the glutamatergic/GABAergic hypothalamic supramammillary nucleus (Hashimotodani et al., 2018) innervate the rILN→DS neurons residing in CM and contralateral CL (Fig. 5C). Previous tracing studies indicate that the rILN receive input from the basal ganglia output projection, the substantia nigra pars reticulata (Kaufman & Rosenquist, 1985), as well as from other targets of nigral output including the superior colliculus (Yamasaki et al., 1986), reticular formation (Krout et al., 2002), and pedunculopontine nucleus (Hallanger et al., 1987; Huerta-Ocampo et al., 2020). We observed that projections from all four of these regions synapse on rILN→DS neurons (Fig. 5D-G).

**Figure 5.**
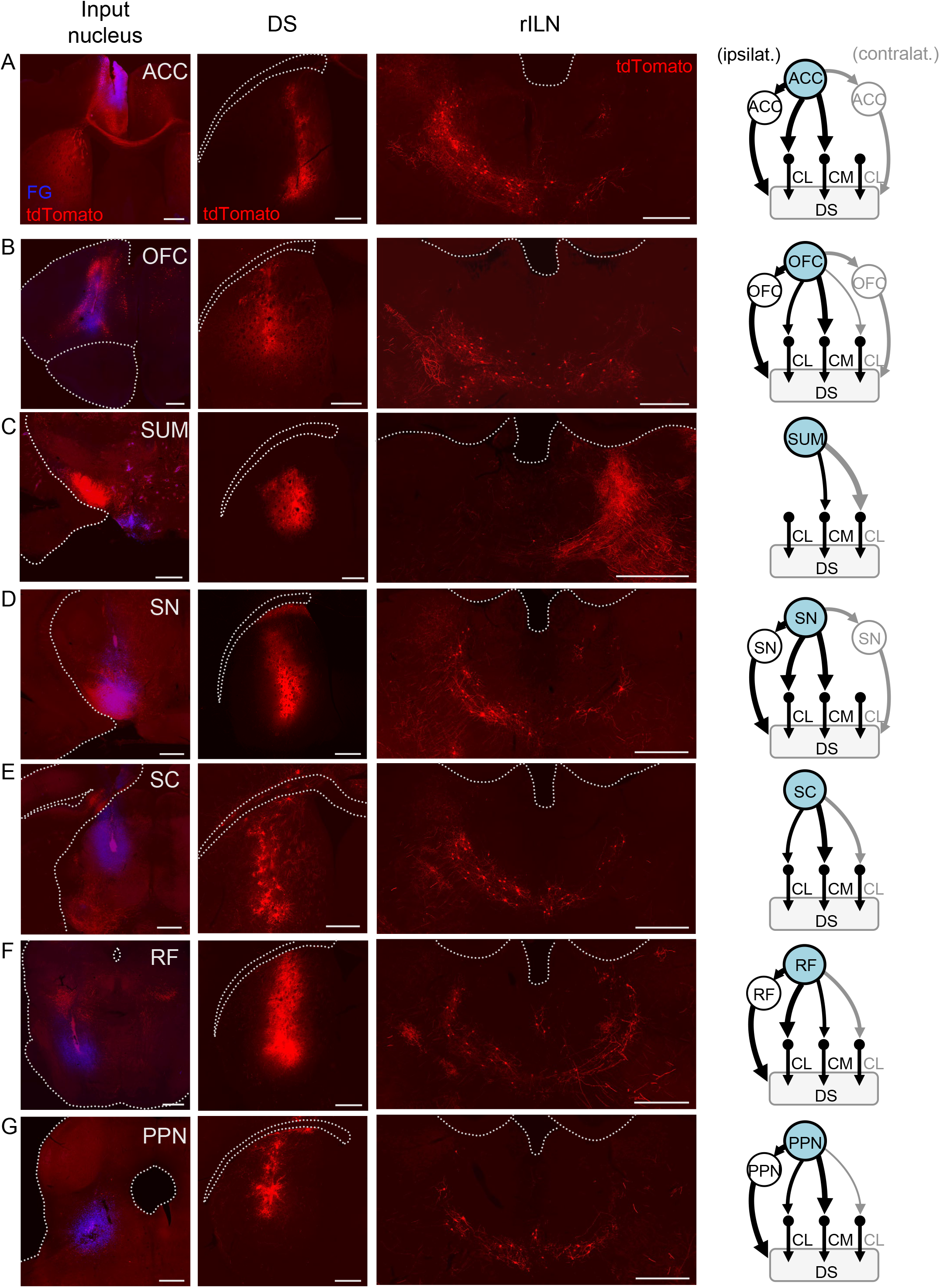
Afferent innervation of rILN→DS neurons. A. Left: injection sites of AAV1-cre and fluorogold (FG) in the anterior cingulate cortex (ACC). Middle left: Injection site for retrograde cre-dependent tdTomato expressing virus in the central DS. Middle right: Representative tdTomato labeling of DS-projecting rILN neurons postsynaptic to ACC projections. Right: Summary of OFC connectivity to rILN nuclei. B-G. Same as A but for orbitofrontal cortex (OFC; B), supramammillary nucleus (SUM; C), substantia nigra (SN; D), superior colliculus (SC; E), reticular formation (RF; F), and pedunculopontine nucleus (PPN; G). Summary diagrams are based on 2-4 cases per region. Scale bars: 500 μm. See also Figure S4.

## Discussion

Our results demonstrate that rILN→DS neurons stably activate at action initiation and reward acquisition and that manipulations of this circuit significantly impact action execution. We found that rILN→DS pathway activation biased mice to initiate more rewarded lever presses whereas inhibitory manipulations resulted in fewer completed press sequences. These results complement our previous findings of rILN suppression reducing overall movement (Cover et al., 2019). Together, these findings implicate the rILN→DS pathway as a critical contributor to action reinforcement and, therefore, performance. We also observed that rILN→DS projection neurons exhibit homogenous physiological properties across the three rILN nuclei, which display relative differences in the afferents impinging upon them. How these differences culminate in unique functional features across these nuclei requires further study.

Corticostriatal circuits guide action learning and habit formation (Kupferschmidt et al., 2017; Smith & Graybiel, 2013; Yin & Knowlton, 2006) by coordinating the activity of striatal output neurons (Tecuapetla et al., 2016; Yin et al., 2009). Thalamic inputs are suggested to influence action expression through modulation of these corticostriatal circuits (Ding et al., 2010). However, rILN signaling also directly drives striatal MSN activity (Chen et al., 2014; Ellender et al., 2013) and cholinergic interneuron firing (Cover et al., 2019). Our observation of rILN→DS neuronal activity occurring at action initiation and correlating to initiation latency suggests that this excitatory input may directly drive striatal signaling for action execution. The absence of discrete signal changes for presses two through five in mice that are well-trained on the FR5 task may reflect the concatenation of the individual movements into a fluid sequence (Jin et al., 2014). Alternatively, press-related rILN→DS neuronal activity may be obscured due to the temporal limits of the calcium sensor and photometric system. Regardless, these results support previous observations of rILN activity mediating motor control (Chen et al., 2014; Giber et al., 2015; Luma et al., 2022) and action switching (Kato et al., 2018).

Our finding that rILN→DS neurons activate at reward acquisition presents, to our knowledge, a unique addition to known reward-related circuitry. Midbrain dopamine neuron firing shifts during reinforcement learning from reward presentation to associated cues and unexpected reward presentation or omission to provide teachable reward prediction errors (Schultz, 1998). In contrast, we found that the rILN stably and persistently signals at reward acquisition across all stages of training and in multiple contexts. Similar reward-related activity is observed in the rILN of primates performing an oculomotor task (Wyder et al., 2003). This signal may serve as a mechanism to provide ongoing reinforcement of appropriately selected actions. Such a mechanism would be advantageous for both the learning of rewarded action sequences but also provides a salient omission when action outcomes change. The sudden absence of this consistent signal may provide a simple cue to drive the search for a new action plan that results in reward. Indeed, we observed a modest negative signal fluctuation when mice prematurely check for sucrose pellets mid-FR5 sequence suggesting that a potential bidirectional signal may provide an instructive cue for action reinforcement. Inhibition of rILN→DS activity significantly reduced the execution of FR5 pressing, which may be due to degradation of this putative reinforcement signal. Regardless, the fiber photometry findings should be considered cautiously given that this approach may not sensitively detect the activity of minority neuronal populations.

Through a cholinergic intermediary, the rILN are capable of locally inducing striatal dopamine release in a behaviorally significant manner (Cover et al., 2019). Our finding of rILN signaling at reward acquisition may suggest that this local dopamine release mechanism is evoked at such events. This possibility is supported by a growing body of evidence identifying discrepancies between dopamine neuron cell body firing and terminal activity. Dopamine release ramps with proximity to rewards and scales to reward magnitude (Hamid et al., 2016; Howe et al., 2013; Mohebi et al., 2019), in addition to dopamine terminal signaling coinciding with reward presentation (Howe & Dombeck, 2016); activity that is not reflected in cell body firing. The rILN may contribute to this phenomenon by locally inducing dopamine release. The rILN may also relay reward-related information directly to MSNs as these cells are shown to encode reward value and action outcome (Hori et al., 2009; Lauwereyns et al., 2002; Nonomura et al., 2018).

One question that emerges from these results is the origin of the rILN→DS action initiation and reward-related signals. Our neuronal tract tracing experiments identified a range of afferents impinging on rILN→DS neurons. The ACC and OFC, which themselves receive input from the rILN (Hunnicutt et al., 2014; Murphy & Deutch, 2018; Van der Werf et al., 2002), provide two sources of excitatory input. These cortical regions are broadly associated with decision making. The OFC encodes outcome value (Gremel & Costa, 2013; Malvaez et al., 2019; Stolyarova & Izquierdo, 2017; Zhou et al., 2019), whereas the ACC encodes decision predictions, surprise signals, and prediction errors, in addition to mediating cognitive control (Hayden et al., 2011; Shenhav et al., 2013; Totah et al., 2009; Wallis & Kennerley, 2011). Although less studied, the SUM is well connected with mesolimbic circuitry and linked to regulation of feeding behavior (Plaisier et al., 2020). Together, these afferents may impart value and motivation for rILN→DS guided action selection.

We also determined that rILN→DS neurons receive input from a basal ganglia output center, the substantia nigra pars reticulata, as well as from nuclei that themselves receive input from the substantia nigra pars reticulata: the superior colliculus, reticular formation, and pedunculopontine nucleus. These results confirm the presence of both direct and indirect basal ganglia subcortical loops relaying back to the DS (Alexander et al., 1986). Through inhibition driven by the substantia nigra and selective disinhibition enabled by D1-MSN pathway signaling, these loops are hypothesized to support the main functions of the basal ganglia: action selection and reinforcement learning (McHaffie et al., 2005; Redgrave et al., 2011). The rILN are uniquely positioned to provide feedback for these functions and our observation of rILN activity corresponding to action initiation and reward acquisition supports this notion. Taken together, rILN pathology may contribute to disorders of action engagement: rILN hyperactivity may contribute to drug seeking (Li et al., 2018; Wang et al., 2019) in a role similar to the adjacent paraventricular thalamus (Hamlin et al., 2009; Matzeu et al., 2015), or to Attention Deficit Hyperactivity Disorder (Jones et al., 2020), while hypoactivity may contribute to impaired goal-directed behavior in Major Depression (Höflich et al., 2019).

## Acknowledgements

The authors thank Dr. Alexandros Poulopoulos for microscopy assistance and Dr. Donna Calu for Pavlovian behavior consultation. BioRender.com was used to create some of the figure schematics. This work was supported by the National Institute on Alcohol Abuse and Alcoholism grant R01AA024845 (to B.N.M.) and National Institute of Drug Abuse grant F31DA047014 (to K.K.C.). The content is solely the responsibility of the authors and does not necessarily represent the official views of the National Institutes of Health.

## Author Contributions

K.K.C. and B.N.M. conceived the experiments. K.K.C. and A.G.L. performed the experiments. K.K.C., A.G.L., and M.M.H. analyzed the experimental data. K.K.C. and B.N.M. wrote the manuscript.

## Materials and Methods

### Subjects

2-5 month-old male and female C57BL/6J (wild-type; Jackson Laboratory, #000664) mice were housed in a temperature and humidity controlled vivarium under a 12-hour light/dark cycle (lights on at 0700 hr). Mice were housed with littermates (2-5) per cage, except for those singly housed following fiber or cannula implantation. Mice performing operant tasks were weighed and fed daily to maintain 90% of ad libitum weight; all others received ad libitum food and water. All experiments were performed in accordance with the United States Public Health Service Guide for Care and Use of Laboratory Animals and were approved by the Institutional Animal Care and Use Committee at the University of Maryland, Baltimore. Sample sizes were determined based on publication standards. Studies were completed across multiple cohorts that produced homogeneous experimental findings. Animals were randomly assigned to control and experimental groups for applicable behavioral experiments. Group assignment was also balanced to ensure equal numbers of males and females between experimental and control cohorts. Blinding was not used in the present study as all subjects and data were treated with identical equipment, analysis methods, and exclusions criteria.

### Surgical Procedures

Mice were anesthetized with isoflurane (5% induction, 1-2% maintenance) and head-fixed in a mouse stereotaxic apparatus (Kopf Instruments, Tujunga, CA). Bupivacaine hydrochloride (0.25%; s.c.) was applied to the scalp prior to incision. Viruses were infused at a rate of 25 nl/min using a 25G syringe (Hamilton Company, Reno, NV). All mice received carprofen (5mg/kg; s.c.) and recovered for a minimum of 7 days to prior to behavioral testing.

For fiber photometry experiments, AAVrg-EF1a-mcherry-IRES-Cre (Addgene #55632-AAVrg) was unilaterally injected in the central DS (600nl; distance from bregma in mm, (AP) +0.65, (ML) 1.70, (DV) −3.00) and two unilateral injections of AAV5-Syn-Flex-GCaMP6s (Addgene #100845-AAV5) were made in the rILN (300 nl/injection; (AP) −1.25 and −1.95, (ML) 0.75, (DV) −3.00). In a separate surgery 3-4 weeks after, a low numerical aperture (NA; 0.22) fiber (200um core, Thorlabs, Newton, NJ) was implanted in the rILN ((AP) −1.60, (ML) 0.76, (DV) −3.00). For optogenetic manipulations, AAVrg-EF1a-mCherry-IRES-Cre was bilaterally injected in the DS (as stated above) and either AAV5-EF1a-DIO-eNpHR3.0-eYFP-WPRE (Addgene #26966-AAV5), AAV5-EF1a-DIO-hSyn-ChR2-eYFP-WPRE (Addgene #20298-AAV5), or AAV5-EF1a-DIO-eYFP-WPRE (Addgene #27056-AAV5) was injected in the rILN (350 nl/hemisphere; (AP) −1.40, (ML) ± 0.75, (DV) −3.50). Optic fibers (NA 0.66, 200um core, Prizmatix, Givat-Shmuel, Israel) were later implanted in the rILN: one at ((AP) −1.60, (ML) −0.75, (DV) −2.70) and the second at a 20° AP angle ((AP) −2.58, (ML) +0.75, (DV) −3.03). All implants were secured with skull screws (BASi, West Lafayette, IN) and dental cement.

For electrophysiology experiments, retrograde AAVrg-CAG-tdTomato (Addgene #59462-AAVrg) was injected in the DS (same coordinates and volume as previously listed) in wild-type mice to patch tdTomato-labeled rILN neurons.

Tran-synaptic tract tracing was performed to identify inputs to rILN→DS neurons. Mice were bilaterally injected with AAVrg-CAG-FLEX-tdTomato-WPRE in the DS (Addgene #28306-AAVrg; 600 nl/hemisphere; (AP) +0.65, (ML) ± 1.70, (DV) −3.00). Afferent regions of interest were unilaterally injected with AAV1-hSyn-Cre-WPRE-hGH (Addgene #105553-AAV1) as well as Fluorogold (20 nl; Fluorochrome LLC, Denver, CO) to label the injection site. The unilateral injection hemisphere was counterbalanced across animals for each input region. Stereotaxic coordinates and virus volumes are as follows: OFC ((AP) +2.50, (ML) 1.00, (DV) −2.20; 250 nl), ACC ((AP) +1.00, (ML) 0.30, (DV) −1.10; 200 nl), SUM ((AP) −2.40, (ML) 0.50, (DV) −5.65; 150 nl), SN ((AP) −3.20, (ML) 1.50, (DV) −4.80; 250 nl), SC ((AP) −3.50, (ML) 0.80, (DV) −2.75; 240 nl), RF ((AP) −4.20, (ML) 0.75, (DV) −4.25; 310 nl), and PPN ((AP) −4.70, (ML) 1.20, (DV) −3.00; 250 nl).

Immunohistochemistry was performed to confirm virus expression and fiber placement; animals were excluded from experiments for poor virus expression at target regions or viral expression in non-target regions (i.e., the caudal intralaminar parafascicular nucleus).

### Immunohistochemistry

Mice were transcardially perfused with room-temperature 0.1M Phosphate Buffered Saline (PBS), pH 7.3, followed by ice-cold 4% (w/v) paraformaldehyde in PBS. Brains were extracted and post-fixed with 4% paraformaldehyde in PBS at 4° C. 50 um coronal sections were cut using a Leica VT100S vibrating microtome. Goat anti-GFP (Abcam #ab6673, Waltham, MA) and chicken anti-mCherry (Novus Biologicals #NBP2-25158, Littleton, CO) primary antibodies were used at a 1:2000 dilution to amplify GCaMP6s and cre expression, respectively. Chicken anti-mCherry was also used to amplify virally-expressed tdTomato. Donkey anti-goat conjugated to Alexa Fluor 488 (Jackson ImmunoResearch #703-545-155, West Grove, PA) and donkey anti-chicken conjugated to Alexa Fluor 594 (Jackson ImmunoResearch #703-585-155) secondary antibodies were used at a 1:1500 dilution. The Brain BLAQ protocol (Kupferschmidt et al., 2015) was used for immunohistochemistry following electrophysiology.

### Slice Electrophysiology

Mice were deeply anesthetized with isoflurane before rapid decapitation and brain removal. 250 μm thick coronal slices were prepared in ice cold, 95% oxygen, 5% carbon dioxide (carbogen)-bubbled modified aCSF (In mM: 194 sucrose, 30 NaCl, 4.5 KCl, 1 MgCl2, 26 NaHCO3, 1.2 NaH2PO4, and 10 D-glucose) before incubating at 32 °C for 30 min in aCSF (in mM: 124 NaCl, 4.5 KCl, 2 CaCl2, 1 MgCl2, 26 NaHCO3, 1.2 NaH2PO4, and 10 D-glucose). Slices were then stored at room temperature until recording.

Thalamic brain slices were transferred to the recording chamber and perfused with carbogen-bubbled aCSF(in mM; 124 NaCl, 4.5 KCl, 2 CaCl2, 1 MgCl2, 26 NaHCO3, 1.2 NaH2PO4, and 10 D-glucose) at 29-31°C. Whole-cell recordings were made with borosilicate glass pipettes (2-5 MΩ). aCSF was perfused on slices through a gravity perfusion system. DS projecting rILN neurons were recorded with a potassium-based internal solution (in mM: 126 potassium gluconate, 4 KCl, 10 HEPES, 4 ATP-Mg, 0.3 GTP-Na, and 10 phosphocreatine; 290-295 mOsm; pH 7.3) containing a hydrazide dye conjugated with Alexa Fluor 488 (40mM) to allow for post-hoc identification of cell location within the rILN. Membrane capacitance and resistance were collected at −60 mV voltage clamp configuration, whereas the resting membrane potential was obtained in current-clamp with a MultiClamp 700B amplifier (Molecular Devices, San Jose, CA); recordings were filtered and digitized at 2 and 10 kHz, respectively. Input resistance was calculated from the mV change induced by a 140 pA hyperpolarizing current step. Data was acquired with Clampex 10.4.1.4 and analyzed with Clampfit 10.4.1.4 (Molecular Devices). Calculation of the accommodation index for rILN AP spiking was based on the entire 0.5 s current injection step that elicited maximum firing as previously calculated (White & Mathur, 2018). Cells that did not maintain sustained firing (i.e. resulted in a depolarization block) were excluded from this analysis. Cells were determined to reside in CL, PC, and CM nuclei based on the Franklin & Paxinos Mouse Atlas (2008).

### Behavioral Testing

#### Fixed-ratio 5 (FR5) lever press operant paradigm

Food-restricted mice were trained to lever press for pellets in an operant chamber (21.6 x 17.8 x 12.7 cm chamber, Med Associates Inc., Fairfax, VT) containing a retractable lever and a trough pellet receptacle equipped with an IR beam that delivered 14 mg sucrose pellets (#F05684, BioServ, Frenchtown, NJ). On each trial, the lever extended into the chamber and retracted once the mouse retrieved a rewarded sucrose pellet (for completed trials) or after the response time limit passes. Mice received two 30-minute training sessions per day with criterion to progress to the next training schedule consisting of ≥ 30 completed fixed-ratio trials for 2 consecutive sessions. Training started on FR1 and progressed to FR3 and FR5, all without time limit restrictions to complete the press sequence. Mice were then trained to complete the FR5 sequence under increasing time constraints (time from lever extension), progressing through 30 s, 15 s, 10 s, 7.5 s, and 5 s schedules. Some mice were unable to successfully perform on FR5-5s and remained at FR5-7.5s for further testing. Optogenetic manipulations were administered once mice achieved consistent performance at or above criterion on their terminal protocol (approximately 8-10 sessions).

A variation of the FR5 lever press task was conducted in which sessions alternated between sucrose pellet reinforcement rates of 100% and (pseudo-randomly) 50% (Fig. S3 H-I). Fiber photometric data was analyzed for all completed FR5 trials following completion of 5 FR5s (for 100% reinforcement sessions) and 5 completed but unreinforced FR5s (50% reinforcement sessions) per session.

#### In vivo optogenetics

For FR5 optogenetic experiments, 470 nm light was delivered bilaterally during experimental sessions using an LED system (Plexon; Dallas, TX). Our group previously demonstrated that 470-nm light sufficiently activates eNpHR channels to inhibit neuronal activity (White et al., 2018, 2020). For optical inhibition, light was delivered pseudo-randomly on 33% of the trials with performance averaged across four consecutive sessions, comparing performance with non-light delivered trials. For optical activation, light was delivered pseudo-randomly on 33% of the trials with performance averaged across six consecutive sessions, comparing performance with non-light delivered trials. For both experiments, mice with the highest and lowest difference in performance were excluded from analysis (both experimental and control cohorts). Experimenters were not blinded to animal group designation.

#### Pavlovian appetitive conditioning

To assess for rILN→DS neuronal activity related to reward-related cues, food-restricted behaviorally naïve mice expressing GCaMP6s in rILN→DS neurons with a multimode fiber implanted in the rILN underwent a Pavlovian conditioned reinforcement paradigm and were randomly assigned to one of two cohorts. For one cohort (“Tone-paired”), a 2 s 10 kHz tone was presented pseudo-randomly during 30 s long trials and co-terminated with the dispensing of a sucrose pellet. A control cohort (“Tone-unpaired”) received tone and pellet each pseudo-randomly. Mice received six 30-minute sessions of these respective paradigms. To test for the effect of reward devaluation on rILN→DS neuronal activity, mice received an additional 3 sessions of their respective protocols that were each preceded by 30 minutes of free feeding of sucrose pellets in their home cage.

#### Movement and Reward Analysis

Mice expressing GCaMP6s in rILN→DS neurons were assessed for movement-related rILN activity in a two-chambered arena (70 x 30 x 25 cm) using Ethovision XT video recording (Noldus, Wageningen, the Netherlands). Following habituation and multiple 15-minute recorded sessions, weigh boats filled with strawberry milk (Nesquik) were placed in two corners of the arena and empty weigh boats were placed in the opposite two corners. rILN→DS photometric activity was aligned to when the center of the mouse body passed within 5 cm of the weigh boats.

#### Sucrose Consumption Assay

To directly correlate rILN→DS photometric activity to reward consumption, mice were placed under a reverse 12-hr light cycle (lights off at 0900) for two weeks before rILN photometry recordings were collected while given access to 2 and 8% (w/v) sucrose water connected to a custom lickometer (Patton et al., 2020). Licks were recorded using Axoscope software (Molecular Devices) and rILN→DS photometry signal was aligned via custom MATLAB code to the start of lick bouts. Bouts were defined as 2 or more licks with an inter-lick interval of less than 2 s.

### Fiber Photometry

Photometry data were collected using a customized in vivo fiber photometry system. Two single-wavelength laser modules were used: a 473 nm laser for optimal GCaMP6s excitation and a 405 nm laser to excite GCaMP6s at its isosbestic wavelength (Opto Engine, Midvale, Utah). Emission from 405 nm excitation was used to control for signal artifacts due to photometry cable motion, background fluorescence, and other sources of noise (Kim et al., 2016). The two lasers were multiplexed at 10 or 15 Hz, resulting in a continuous 20 or 30 Hz pulse train. Both laser beams were bounced into a dichroic filter cube designed for 473 and 405 nm excitation as well as for 510 nm emission (Chroma Technology Corp., Bellow Falls, VT). The two excitation wavelengths were focused through a 4x fluorite objective (Olympus, Tokyo, Japan) onto a multimode fiber bundle (Thorlabs). One fiber was connected to the mouse through a chronic unilateral multimode fiber implant and another fiber was placed inside a tube of Alexa Fluor 488 to control for variability in laser energy. Emissions from GCaMP6s and Alexa Fluor 488 were detected as an image of the fiber bundle using an ORCA-Flash4.0 LT high-resolution CMOS camera (Hamamatsu Photonics, Hamamatsu City, Japan). Laser multiplexing and image acquisition were synchronized using an Arduino Leonardo microcontroller. Trial or time-dependent recordings were initiated through MedPC or Ethovision systems. Camera image acquisition parameters were controlled through HCImage software for Hamamatsu cameras.

### Quantification and Statistical Analysis

Electrophysiological data were analyzed using Clampex software. Statistical analyses were performed in Prism (version 6.01; GraphPad Software, San Diego, CA) or MATLAB (R2019a; The MathWorks, Inc., Natick, MA).

Photometry data were analyzed using a combination of custom MATLAB code and Prism. Pixel intensities imaged from the fiber implanted in the rILN and the fluorophore control fiber were first averaged. The background signal in the absence of laser transmission was then subtracted from the averaged signals. Two separate regressions were performed to minimize any noise sources. First, 473 and 405 nm photometry signals were regressed with the corresponding control fluorophore signal, and the residuals of the regression were then used for further processing. The 473 nm signal was then regressed with the 405 nm signal as a covariate, and the residuals of the regression were extracted as the fully processed photometry signal (z-score) from the rILN (White et al., 2020). The photometry signal was then statistically analyzed based on performance or operant task event (i.e., movement-aligned activity or FR trial outcome). For each analysis comparing two or more categories of rILN signal, the number of averaged events or trials were standardized across each compared category for each animal. The average z-score for three consecutive time points were statistically analyzed with the appropriate t-test or ANOVA. Signal area under the curve was computed in MATLAB.

To quantify relative strength of afferent innervation to rILN→DS neurons identified in TranSTarT cases, tdTomato-labeled cells were counted in CM, ipsilateral CL, and contralateral CL. Separate ratios of cell counts for CM: ipsilateral CL and ipsilateral CL: contralateral CL were derived, correcting for area, within each coronal slice and averaged across slices and cases for a given assessed afferent. Ratios indicating a difference ≥ 40% of labeled neurons between the compared rILN nuclei are denoted by line thickness in Figure 5 summary diagrams. This criterion was also used to assess relative strength of ipsilateral and contralateral afferent→afferent→DS di-synaptic circuits, when applicable.

All statistical analyses are reported in the Results. The specific statistical test as well as value and description of n are listed in the figure legends. Averaged data is expressed as mean ± standard error. Correlations are expressed as Pearson’s r and 95% confidence intervals. All statistical tests were conducted two-sided, when applicable. Significant post-hoc Holm-Ŝidàk tests for ANOVAs are indicated in the figures. All asterisks in figures indicate: * P < 0.05, ** P < 0.01, *** P < 0.001, **** P < 0.0001.

### Data Availability

All data generated or analysed during this study are included in the manuscript and supporting files; Source Data files have been provided for all Figures.

---

**Supplemental Figure S1.**
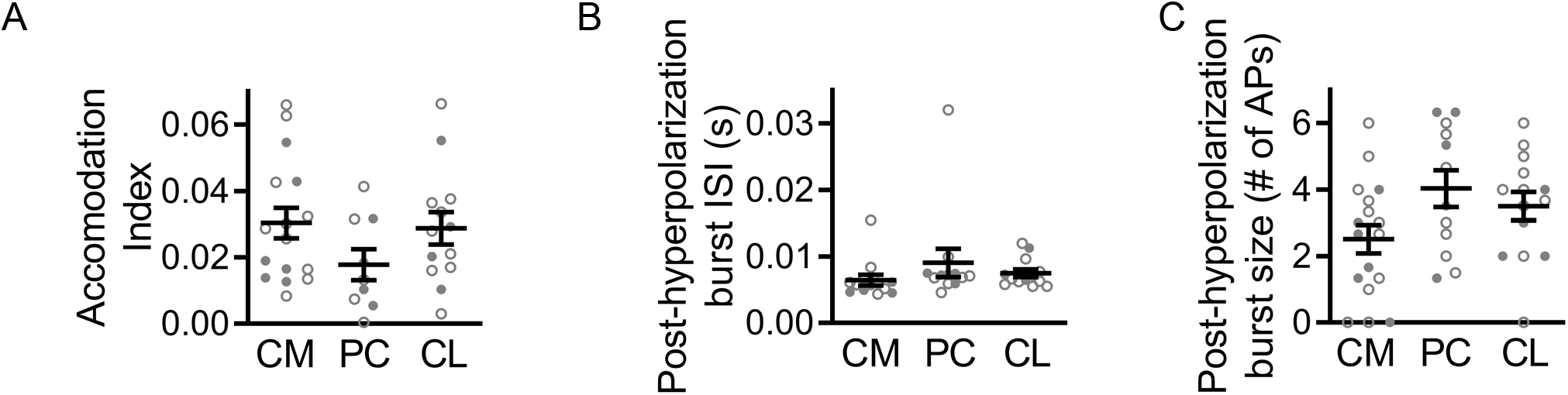
Additional rILN firing properties. A. The accommodation index for the maximum rate of action potential (AP) firing in response to a depolarizing current step did not differ significantly between CM (n=16), PC (n=9), and CL (n=12) neurons. B. AP burst firing following hyperpolarizing current steps did not significantly differ in inter-spike interval (ISI) between CM (n=13), PC (n=12), and CL (n=12) neurons. C. The number of APs evoked following hyperpolarizing current steps did not differ significantly between CM (n=17), PC (n=12), and CL (n=13) neurons. One-way ANOVA (A-C). n = number of cells. Filled and open data points represent cells from male and female mice, respectively.

**Supplemental Figure S2.**
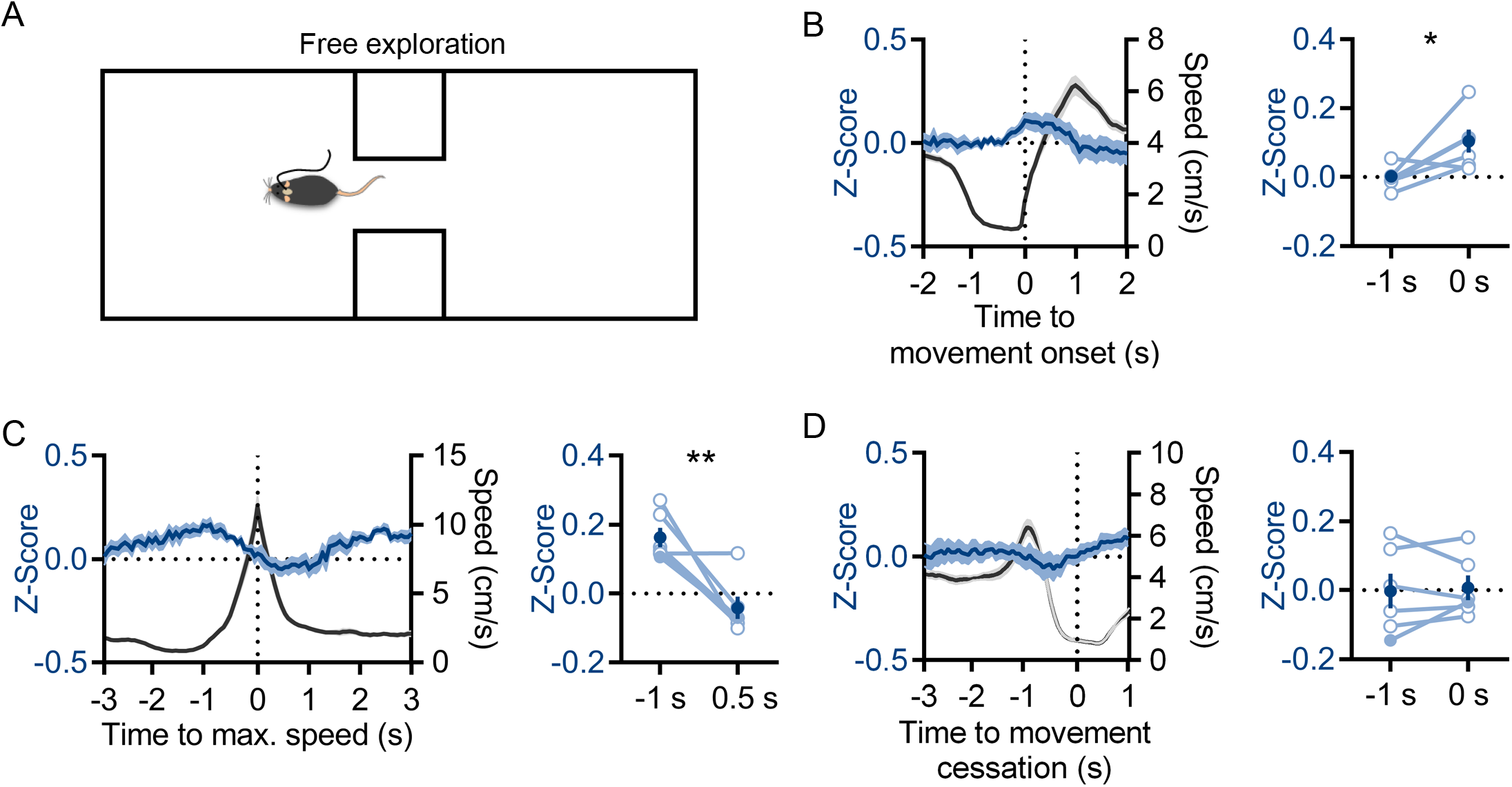
rILN→DS neuronal activity correlates with movement initiation. A. Movement speed-aligned rILN→DS activity was recorded in mice freely moving in an open arena. B. Left: average rILN→DS activity (blue) aligned to movement onset (gray). Right: rILN→DS activity increased at the time of movement onset (N=6). C. Left: rILN→DS activity aligned to maximum speed in movement epochs. Right: rILN→ signal peak proceeded movement epochs. D. Left: rILN→DS activity aligned to movement cessation. Right: Cessation did not correspond to changes in rILN→DS activity. Paired t-test (B-D). N = number of mice. Filled and open data points represent male and female mice, respectively.

**Supplemental Figure S3.**
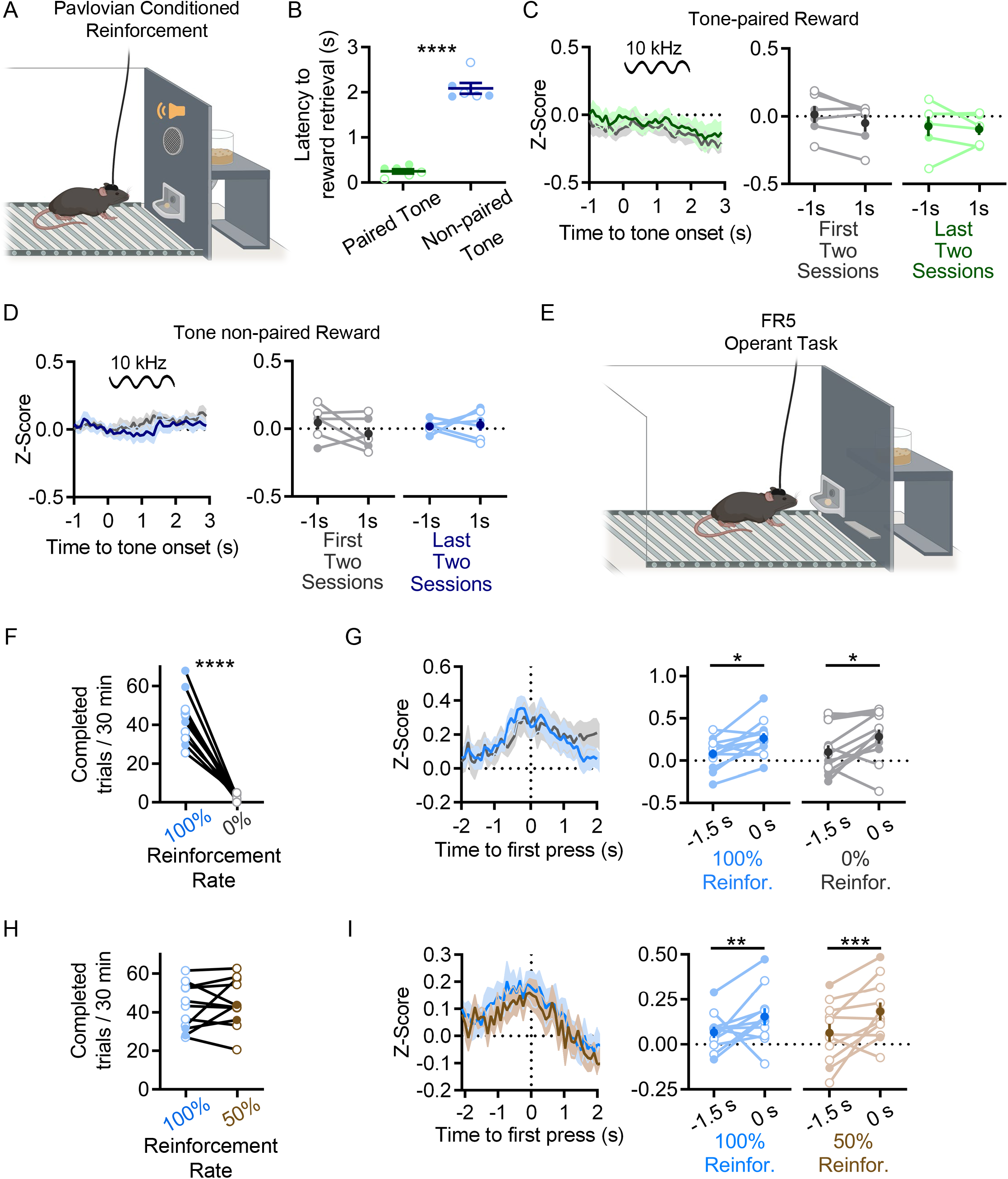
rILN→DS activity does not correlate with reward-predicting Pavlovian cues or changes in operant task reward probability. A. Mice underwent a Pavlovian conditioning paradigm in which sucrose pellet delivery immediately followed a pseudo-randomly presented 2 s long tone (tone paired) or in the control group, 2 s tones and sucrose pellets were pseudo-randomly delivered (tone non-paired). B. In the last two conditioning sessions, the average latency to retrieve delivered sucrose pellets was significantly shorter for the tone-paired group than the tone non-paired control cohort (t=14.46, P<0.0001). C. Fiber photometric recordings of rILN→DS neurons did not show significant differences in activity 1 s before and 1 s after the onset of the tone in either early (gray) or late (green) conditioning sessions (N=6). D. rILN→DS activity in the non-paired cohort was not significantly different 1 s before and 1 s after the onset of the tone in either early (gray) or late (blue) conditioning sessions (N=6). E. Reinforcement manipulations were conducted in mice well-trained on the FR5 lever press task in F-I. F-G. rILN→DS activity was recorded from mice in which completed FR5s were reinforced (blue) or not reinforced (gray) with sucrose pellets in separate sessions (N=12). F. Mice extinguished FR5 responding over 2-3 sessions in which correct responses were not reinforced (data shown from last session). G. Left: First press-aligned rILN→DS activity on completed FR5 trials from reinforced (blue) or unreinforced (gray) sessions. Right: Comparison of rILN→DS signal at −1.5 s and 0 s relative to first press on completed FR5 trials. H-I. rILN→DS activity was recorded in a separate cohort of mice that alternated between FR5 sessions in which completed trials were reinforced at 100% (blue) or 50% (brown) rates (N=11). H. The number of completed FR5 trials did not significantly differ between 100% and 50% reinforcement rate sessions. I. Left: First press-aligned rILN activity on completed FR5 trials for each reinforcement rate. Right: Comparison of average rILN→DS activity at −1.5 s and 0 s relative to first press on completed trials at each reinforcement rate. Unpaired t-test (B), two-way repeated measures ANOVA (C-D, G, I), paired t-test (F, H). N = number of mice. Filled and open data points represent male and female mice, respectively.

**Supplemental Figure S4.**
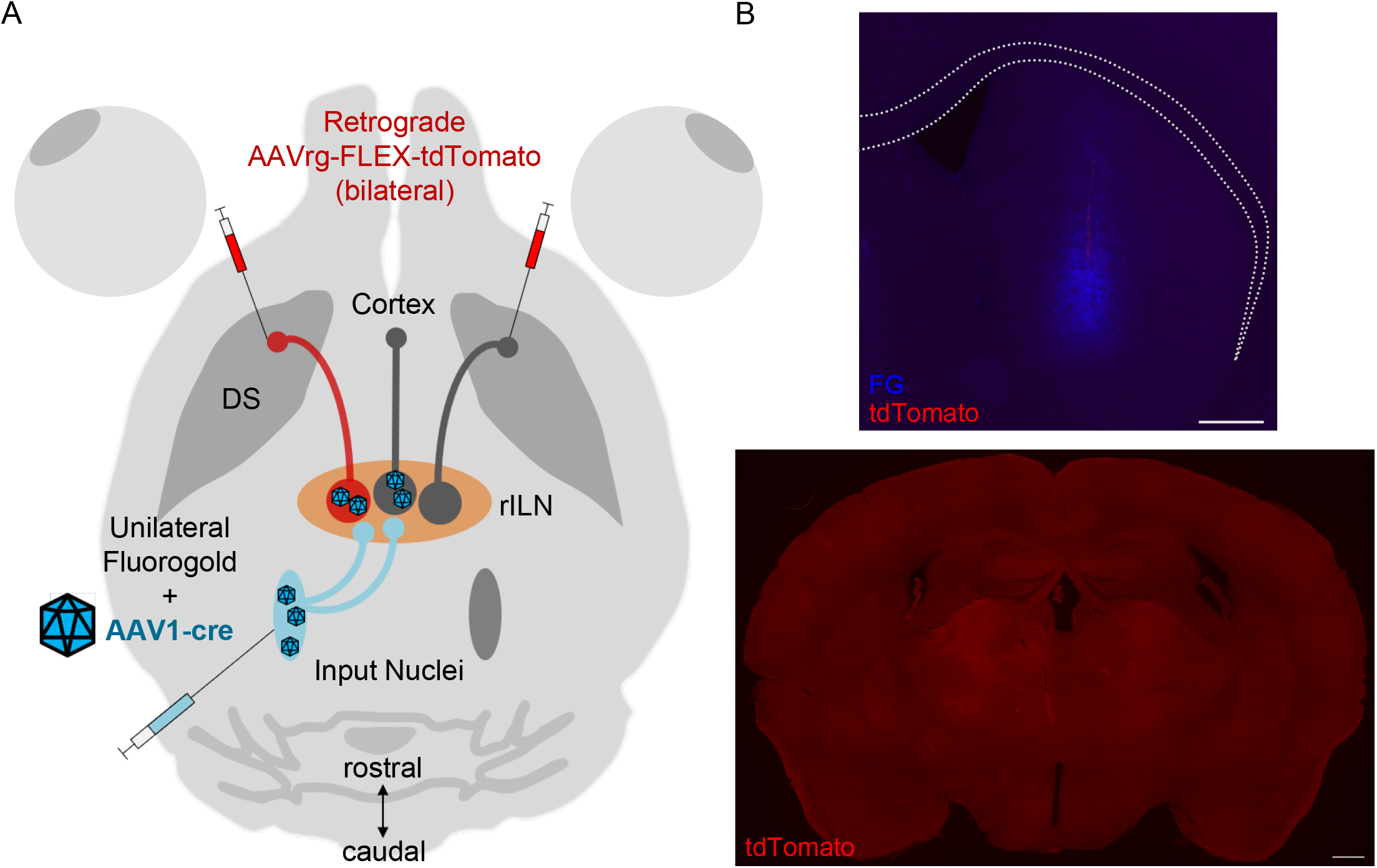
Tran-synaptic neuronal tract tracing to identify specific afferents to the rILN→DS circuit. A. Experimental strategy. An anterograde trans-synaptic virus expressing cre recombinase (AAV1-cre) was unilaterally injected into a nucleus that projects to the rILN, while a retrograde virus that cre-dependently expresses tdTomato (AAVrg-FLEX-tdTomato) was bilaterally injected into the DS. B. Top: Injection site for fluorogold (FG) (as an injection site marker) and AAVrg-FLEX-tdTomato in DS. Bottom: image of thalamus showing no tdTomato labeling in the absence of AAV1-cre injection; demonstrating the necessity for viral transfection of cre for tdTomato expression in efferent cells. Scale bars: 500 μm.

**Supplemental Table S1.** rILN→DS manipulation alters FR5 task performance. Group means and SEM are reported for experimental and control cohorts in percent completed (left), omitted (middle), and incomplete (right) trials, as related to Figure 4 C-E (top) and H-J (bottom).

|  | % Completed trials |  | % Omitted trials |  | % Incomplete trials |  |
| --- | --- | --- | --- | --- | --- | --- |
| *Optogenetic inhibition* | *No light* | *Light* | *No light* | *Light* | *No light* | *Light* |
| eYFP | 42.0 ± 2.2 | 38.7 ± 2.7 | 38.3 ± 2.4 | 38.4 ± 2.3 | 19.7 ± 1.4 | 22.9 ± 2.0 |
| NpHR-eYFP | 41.6 ± 2.7 | 34.2 ± 3.6 | 33.7 ± 3.4 | 39.9 ± 3.7 | 24.7 ± 2.1 | 25.9 ± 2.2 |
| *Optogenetic activation* | *No light* | *Light* | *No light* | *Light* | *No light* | *Light* |
| eYFP | 38.6 ± 2.5 | 40.7 ± 2.4 | 37.6 ± 2.5 | 36.2 ± 2.4 | 23.8 ± 1.7 | 23.2 ± 1.8 |
| ChR2-eYFP | 32.6 ± 2.1 | 46.1 ± 1.9 | 41.3 ± 3.0 | 27.8 ± 2.2 | 26.1 ± 2.0 | 25.8 ± 1.5 |

